# The parasubthalamic nucleus refeeding ensemble delays feeding initiation

**DOI:** 10.1101/2023.01.28.525750

**Authors:** Jeffery L Dunning, Catherine Lopez, Colton Krull, Max Kreifeldt, Maggie Angelo, Charu Ramakrishnan, Karl Deisseroth, Candice Contet

## Abstract

The parasubthalamic nucleus (PSTN) is responsive to refeeding after food deprivation and PSTN subpopulations can suppress feeding. However, no study directly addressed the role of PSTN neurons activated upon food access resumption. Here we show that the ensemble of refeeding-activated PSTN neurons drastically increases the latency to initiate refeeding with both familiar and novel food but exerts limited control over the amount of food consumed by hungry mice. This ensemble also delays sucrose consumption but accelerates water consumption in thirsty mice. We next sought to identify which subpopulations of PSTN neurons might be driving these effects. We discovered that PSTN *Tac1* neurons projecting to the CeA selectively suppress feeding initiation while PSTN *Crh* neurons surprisingly promote the consumption of novel, palatable substances. Our results demonstrate the key role of endogenous PSTN activity in the control of feeding initiation and identify PSTN subpopulations counteracting each other’s influence on consummatory behaviors.

## Introduction

The hypothalamus is a critical hub for the integration of interoceptive signals and the initiation of homeostatic behaviors regulating need states such as hunger and thirst. In particular, molecularly defined subpopulations of neurons in the arcuate nucleus, paraventricular nucleus, and lateral hypothalamic area control food intake (1, 2), while median preoptic nucleus excitatory and inhibitory neurons promote and suppress, respectively, water drinking (3, 4). Beyond physiological needs, hypothalamic nuclei also control food and fluid consumption driven by hedonic motives (i.e., pleasurable sensory perception or affective state), as they interface with corticolimbic and midbrain monoaminergic systems that encode reward and decision making (5, 6).

In recent years, the parasubthalamic nucleus (PSTN), a differentiation of the premammillary lateral hypothalamic area that is highly interconnected within brain regions regulating interoception and appetite, has emerged as a new player in the regulation of ingestive behaviors (see 7 for review). Significant induction of c-Fos expression, a marker of neuronal activation, has been observed in the rodent PSTN following predatory hunting of cockroaches, early active phase surge of food intake, refeeding after food deprivation, anorexia-inducing amino-acid deficient diet refeeding, and sucrose drinking following water deprivation (8–14). While all PSTN neurons are glutamatergic (VGluT2-positive), they comprise subpopulations expressing high levels of either *Tac1* (encoding preprotackykinin-A, a precursor of substance P and neurokinin A) or *Crh* (encoding corticotropin-releasing factor, CRF) with minimal overlap (14–16). Both PSTN^*Tac1*^ and PSTN^*Crh*^ neurons show strong activation in response to refeeding in food-deprived mice (14).

Manipulations of PSTN neurons via optogenetic, chemogenetic, and targeted cell ablation approaches have further demonstrated their functional implication in consummatory behaviors (see 7 for review). Specifically, stimulation of PSTN^VGluT2^ neurons projecting to the paraventricular nucleus of the thalamus (PVT), PSTN^*Tac1*^ neurons as a whole, as well as PSTN^*Tac1*^ neurons projecting to the central nucleus of the amygdala (CeA), PVT, parabrachial nucleus, or nucleus of the tractus solitarius, reduces food intake in *ad libitum* fed mice (14, 17). This effect is not observed when stimulating PSTN^*Crh*^ neurons or PSTN^*Tac1*^ neurons projecting to the bed nucleus of the stria terminalis (BNST), highlighting the cell-type and pathway specificity of the PSTN’s influence on food intake (14). On the other hand, inhibiting PSTN^*Tac1*^ somas counters the anorectic effect of a malaise-inducing agent (lypopolysaccharide), neophobia (first-time exposure to sucrose), or appetite-suppressing hormones (amylin, cholecystokinin, peptide YY), while ablating PSTN^*VGluT2*^ neurons abolishes anorexia induced by glucagon-like peptide-1 (12, 14, 18).

While these studies elegantly demonstrated the ability of several PSTN subpopulations to suppress feeding, none of them directly addressed the role of PSTN neurons activated upon food access resumption in hungry animals in the control of ensuing food consumption. To address this gap of knowledge, we used chemogenetics in “Targeted Recombination in Active Populations” mice (TRAP2 mice, in which the sequence encoding iCre-ER^T2^ is inserted in the *Fos* locus without disrupting endogenous *Fos* expression (19)), to selectively re-activate the ensemble of refeeding-activated PSTN neurons and determine their influence on consummatory behaviors.

We found that, in hungry mice, the PSTN refeeding ensemble drastically increases the latency to initiate refeeding with both familiar chow and a novel, palatable food (Froot Loops) but exerts limited control over the amount of food consumed. In thirsty mice, this ensemble also delays sucrose consumption but accelerates water consumption, with again no influence on the amount of fluid consumed. Given that previous studies only reported measures of intake (i.e., amount consumed), we next sought to examine which subpopulations of PSTN neurons might be driving these latency effects, using cell-type and pathway-specific chemogenetic manipulations.

## Methods

### Animals

TRAP2 (Fos^tm2.1(icre/ERT2)Luo/J^, stock #030323 (19)), *Crh*-Cre (*Crh*-IRES-Cre, B6(Cg)-Crh^tm1(cre)Zjh/J^, stock #012704 (20)), and *Tac1*-Cre (B6;129S-Tac1^tm1.1(cre)Hze/J^, stock #021877 (21)), breeders were obtained from The Jackson Laboratory. C57BL/6J mice were obtained from Scripps Research rodent breeding colony. All mice used for experimentation were heterozygous for the Cre allele. All experimental cohorts contained males and females, and mice were matched by sex and age across experimental subgroups.

Mice were maintained on a 12 h/12 h light/dark cycle. Food (Teklad LM-485, Envigo) and reverse osmosis purified water were available *ad libitum*, except for a 24-hour period of food deprivation or 4-hour water deprivation in relevant experiments. All mice were single-housed in static caging with Sani-Chips (Envigo) bedding one week prior to behavioral assays and remained in these housing conditions for the duration of experimentation. All mice were at least 10 weeks old at the time of surgery. All procedures adhered to the National Institutes of Health Guide for the Care and Use of Laboratory Animals and were approved by the Institutional Animal Care and Use Committee of The Scripps Research Institute.

### Drugs

For iCre-ER^T2^ activation in TRAP2 mice (19, 22), the estrogen receptor ligand 4-hydroxytamoxifen (4-OHT) was obtained from Hello Bio (HB6040). 4-OHT was dissolved in 100% ethanol and then mixed 1:1 with a mixture of 1 part castor oil (Sigma-Aldrich 259853) and 4 parts sunflower oil (Sigma-Aldrich S5007). Ethanol was then removed via vacuum centrifugation and the remaining oil was again diluted in the same 1:4 part oil mixture to achieve a concentration of 10 mg/mL before intraperitoneal (i.p.) injection at a dose of 50 mg/kg (5 mL/kg injection volume, 23 gauge needle). Clozapine-N-oxide (CNO) was used as the ligand for hM3Dq and hM4Di designer receptors (23, 24) and was obtained from Enzo Life Sciences Inc. (BML-NS105-0025). CNO was dissolved in dimethyl sulfoxide (DMSO) and diluted in 0.9% saline (0.5% final DMSO concentration) for i.p. injection (3 mg/kg body weight, 10 mL/kg injection volume, 27 gauge needle). Salvinorin B (SalB) was used as the ligand for the κ-opioid receptor-based designer receptor (KORD (25)) and was obtained from Hello Bio (HB4887). SalB was dissolved in 100% DMSO for subcutaneous (s.c.) injection (10 mg/kg body weight, 1 mL/kg injection volume using a Hamilton 250 μL syringe #81108, 27 gauge, point style 4 needle). Prior to SalB testing, mice were habituated for two days to DMSO s.c. injections.

### Viral vectors

Adeno-associated viral serotype 2 (AAV2) vectors encoding the hM3Dq excitatory or hM4Di inhibitory designer receptor fused to the red fluorescent protein mCherry, or mCherry alone, under the control of the human synapsin promoter (hSyn) and in a Cre-dependent manner (Double-floxed Inverted Open reading frame, DIO), were obtained from the Vector Core at the University of North Carolina at Chapel Hill (AAV2-hSyn-DIO-hM3Dq-mCherry, Addgene plasmid # 44361, lot 8269, titer 1.5 x 10^13^ vg/mL; AAV2-hSyn-DIO-hM4Di-mCherry, Addgene plasmid # 44362, lot 8268, titer 1.8 X 10^13^ vg/mL; AAV2-hSyn-DIO-mCherry, Addgene plasmid # 50459, lot 8267, titer 1.3 x 10^13^ vg/mL) (24). An AAV8 encoding the inhibitory designer receptor KORD fused to the fluorescent protein mCitrine under the control of the human synapsin promoter in a Cre-dependent manner was obtained from Addgene (AAV8-hSyn-dF-HA-KORD-IRES-mCitrine, plasmid # 65417, lot v43122, titer 2.1 X 10^13^ gc/mL) (25). A retrograde AAV (26) encoding the Cre enzyme fused to the green fluorescent protein (GFP) under the control of the human synapsin promoter was obtained from Addgene (AAVrg.hSyn.HI.eGFP-Cre.WPRE.SV40, plasmid # 105540, lot V102961, titer 2.5 X 10^13^ gc/mL). An AAV8 vector expressing hM3Dq-mCherry under a short EF1α promoter in a Cre- and Flp- dependent manner was generated, packaged, and purified by the laboratory of Karl Deisseroth (AAV8-nEF-Con/Fon-hM3Dq-mCherry, lot 7280, titer 1.23 X 10^12^ gc/mL) and used in conjunction with a retrograde AAV encoding the Flpo enzyme under an EF1α promoter (AAVrg-EF1a-Flpo, Addgene plasmid # 55637, lot v56725, titer 1.02 X 10^12^ gc/mL) (27).

### Experimental cohorts

The data were collected from five separate cohorts of mice.

A cohort of 16 TRAP2 mice (7 males + 9 females) featured three experimental subgroups: food-deprived mice injected with 4-OHT (n=6), food-deprived mice injected with vehicle (to control for leaky Cre activity, n=5), and *ad libitum* fed mice injected with 4-OHT (to control for baseline PSTN activity, n=5). In this experiment, all mice were injected with the hM3Dq vector and the effect of CNO was tested according to a between-subject design. For the induction of Cre activity, all mice were transferred to a clean, new cage (with or without food), and 24 hours later, 4-OHT (or vehicle) was injected immediately prior to the placement of chow pellets in the wire lid of food-deprived cages. Deprivation-induced body weight loss was confirmed and vigorous interactions of refed mice with the food hopper were noted, but consumption measures were not collected to avoid interfering with the cages during the time window of Cre activation.

A cohort of 46 *Tac1-*Cre mice (22 males + 24 females) featured three experimental subgroups injected with either the hM3Dq (n=16), hM4Di (n=13), or mCherry (n=15) vectors. The effect of CNO was tested according to a between-subjects design.

A cohort of 19 *Crh*-Cre mice (10 males + 9 females) were all injected with the hM3Dq vector and were treated with either CNO or vehicle for a between-subject analysis. The treatment assigned to each mouse remained the same across all assays.

A cohort of 12 C57BL/6J mice (8 males + 4 females) featured two experimental subgroups that were co-injected with the hM3Dq and KORD vectors (1:1 premixed cocktail) in the PSTN and the retrograde Cre vector in the CeA (n=6) or the BNST (n=6) for pathway-specific manipulations. In this experiment, all mice were injected with chemogenetic ligands (CNO or SalB) and their respective vehicle for within-subject analysis (i.e., each mouse was tested four times). CNO was tested first, then SalB. In each case, the order of ligand and vehicle administration was counterbalanced between mice.

A cohort of 6 *Tac1-*Cre mice (1 male + 5 females) and 7 *Crh*-Cre mice (3 males + 4 females) were injected with the Con/Fon hM3Dq vector into the PSTN and retrograde Flpo vector in the CeA. All mice were injected with CNO and vehicle for within-subject analysis.

### Stereotaxic surgery

All mice were anesthetized with isoflurane and placed in a stereotaxic frame (David Kopf Instruments, model 940). A small hole was drilled in the skull (David Kopf Instruments, 1474) and a 75-200 nL volume of viral vector was injected using a microinjector pump (World Precision Instruments, UMP3T-2) with an attached 10-μL NanoFil syringe (World Precision Instruments) fitted with a 33-gauge NanoFil blunted tip (World Precision Instruments, NF33BL). The following coordinates were used (AP from bregma, ML from midline, DV from skull, in mm): PSTN (AP −2.3, ML ± 1.1, DV −5.2), CeA (AP −1.2, ML ± 2.8, DV −4.5), BNST (AP 0.2, ML ± 1.0, DV −4.3). The vector was infused at a rate of 0.1 μL/min with the needle left in place for 5 minutes post injection to minimize backflow. The scalp was sutured using surgical thread. Mice were left undisturbed for at least three weeks post-injection prior to behavioral testing.

### Histology

Mice were anesthetized with chloral hydrate and perfused with cold phosphate buffered saline (PBS) followed by 3.7% paraformaldehyde (PFA). Brains were dissected and immersion fixed in PFA for 2 hours at 4°C, cryoprotected in 30% sucrose in PBS at 4°C until brains sank, flash frozen in 2-methyl-butane chilled on a dry ice ethanol slurry and stored at −80°C. Coronal 35-μm thick brain sections were sliced with a cryostat (Leica CM1950), collected in five series spanning appropriate brain regions in PBS containing 0.01% sodium azide, and stored at 4°C. Sections were then washed for 10 minutes in PBS, plated on Superfrost plus glass slides (Fisher Scientific, 1255015), and air-dried. Coverslips were mounted using DAPI-containing Vectashield Hardset medium (Vector Laboratories, H1500). Images were captured using a Keyence BZ-X700 fluorescence microscope. In all DIO cohorts, native fluorescence signals were used to assess the accuracy of stereotaxic targeting and exclude mistargeted mice from behavioral datasets accordingly (sample sizes reported in the “Experimental cohorts” section include well-targeted mice only). In pathway-targeted mice, we were not able to image mCitrine native fluorescence due to overlapping signal from eGFP-Cre in the green channel but used the mCherry signal to assess the location of the KORD/hM3Dq cocktail infusion. In the Con/Fon cohort, we used mCherry immunolabeling to enhance the signal (chicken anti-mCherry antibody, Abcam, ab205402, RRID AB_2722769, 1:5,000, overnight incubation at 4°C; goat anti-chicken conjugated to Alexa Fluor 568, Life Technologies, RRID AB_2534098, 1:500, 2-h incubation at room temperature). Counts of left and right hemisphere mCherry-labeled cells in TRAP2 mice were obtained with the Cell Counter tool of ImageJ (NIH), averaged, and used to evaluate the size of the refeeding-activated PSTN ensemble.

### Behavioral testing

All behavioral testing was conducted during the active (dark) phase of the circadian cycle, 3-6 hours after lights had turned off. For all assays, CNO, SalB, or their respective vehicle was administered 30 min prior to the introduction of food/fluid. Latency to first bite/lick was recorded as a measure of motivation to consume, and the amount of food/fluid consumed was measured over 30 min (except for water consumption in TRAP2 mice, which was measured over 2 hours).

### Testing with solid foods

A single pre-weighed food pellet (familiar chow) or seven pieces of Froot Loop cereal (novel or habituated) were introduced in the cage. The wire lid, water bottle, and filter top were then put back in place. Latency to first bite was collected with a stopwatch, with the experimenter carefully distinguishing between movement of the food, burying of the food, and actual biting of the food which was accompanied by mastication sounds. The chow pellet or Froot Loops were weighed again 30 minutes later to measure amount of food consumed.

For food deprivation, mice were transferred to a new, clean home cage without food and testing was started 24 hours later.

Preference testing for chow versus Froot Loops was performed in *Tac1*-Cre mice, in the absence of chemogenetic manipulations. *Ad libitum* fed mice were given two chow pellets and 20 pieces of Froot Loop cereal that were measured before and after a 24-hour period. 19 of 46 mice consumed all the Froot Loops during the testing period, the calculated preference values are therefore an underestimate of actual preference.

For Froot Loops habituation, mice were given chow and Froot Loops *ad libitum* for another three days after preference testing.

For experiments in sated *Tac1*-Cre mice, mice were given scheduled access to three pieces of Froot Loops 3 hours into the dark phase for 7 or 8 days prior to the experiment, along with *ad libitum* access to chow.

For SalB testing in pathway-targeted mice, mice were tested in an open arena (Taconic Transit Cage, 56-cm long × 40-cm wide × 18-cm deep) lined with 2 cm of fresh bedding, instead of their home cage. A single food pellet was secured onto a platform located in the center of the arena (see 28 for details). This novel environment was used to make the mice more hesitant to initiate feeding and facilitate the detection of a latency reduction.

### Testing with liquids

Prior to liquid testing, mice were habituated to 50-mL conical tubes fitted with rubber stoppers and sippers for 7 days. On the morning of testing, water bottles in each cage were removed at the onset of the dark phase for 4 hours prior to testing. Mice consume very little water during their inactive phase and a significant portion of their daily water intake at the beginning of their active phase (29, 30), such that mice deprived of water for the first 4 hours after lights turn off have a high motivation to consume water when access is resumed. At the time of testing, a single, pre-weighed 50-mL conical tube containing normal drinking water or a 5% (w/v) solution of sucrose (Sigma-Aldrich 84097) was inserted in the wire lid, and the filter top was put back in place. Latency to first lick was collected with a stopwatch, with attention paid to differentiating whisking from licking behaviors. A 5-minute time limit was used for latency measurements. The bottle was weighed again 30 minutes (or 120 minutes for water testing in TRAP2 mice) later. A control cage with no mouse was used as spill control and the weight lost in that cage was subtracted from all other cages.

Preference testing for water versus sucrose was performed in *Tac1*-Cre mice, in the absence of chemogenetic manipulations. All mice were given two 50-mL conical tubes in their home cage for 72 hours, one containing water and one containing 5% sucrose solution. Bottles were weighed before and after the experiment. The 72-hour preference test was used to habituate mice to the sucrose solution for subsequent experiments.

### Statistics

Raw data was processed in Microsoft Excel and statistical analysis was performed using GraphPad Prism software. Experiments evaluating between-subject effects of CNO in TRAP2 (including mCherry counts) and *Tac1*-Cre cohorts were analyzed using ordinary one-way ANOVAs, with an alpha of 0.05. Multiple comparisons were conducted with the Tukey’s test for TRAP2 mice and with the Dunnett’s test for *Tac1*-Cre mice (using the mCherry group as control condition). Experiments comparing the effect of CNO versus vehicle were analyzed using either unpaired (*Crh*-Cre mice) or paired (pathway-targeted and Con/Fon cohorts) t-tests, with an alpha of 0.05. In each graph, individual values are plotted, bars show group averages, and error bars represent standard error of the mean.

## Results

### The PSTN refeeding ensemble delays the latency to initiate feeding

Subpopulations of PSTN neurons become highly active in response to refeeding after food deprivation, as indexed by c-Fos induction (10, 13, 14). To determine the functional significance of this neuronal ensemble, TRAP2 mice were injected with a Cre-dependent hM3Dq-encoding virus in the PSTN and administered 4-OHT upon refeeding following 24 hours of food deprivation. In addition to this experimental group, a first control group was injected with 4-OHT in a sated state to control for baseline PSTN activity, and another control group was injected with vehicle upon refeeding to control for leaky (i.e., 4-OHT-independent) Cre recombination (**Figure 1A**). Native fluorescence of the mCherry reporter was used to quantify the size of the PSTN neuronal ensemble targeted in each of the 3 experimental groups. As expected, there were very few mCherry labeled cells in vehicle-injected mice (**Figure 1B**). Accordingly, there was a significant main effect of group (F(2,12)=14.68, p=0.0006), whereby mice injected with 4-OHT at the time of refeeding (p=0.0004) and those injected with 4-OHT in a sated state (p=0.0447) had significantly more mCherry positive cells than vehicle-injected controls. The ensemble captured in a sated state, which reflects baseline PSTN activity, was significantly smaller than that captured upon refeeding (p=0.0482) but provides a sizeable population of PSTN cells to inform on the functional specificity of the PSTN refeeding ensemble upon chemogenetic manipulation.

**FIGURE 1.**
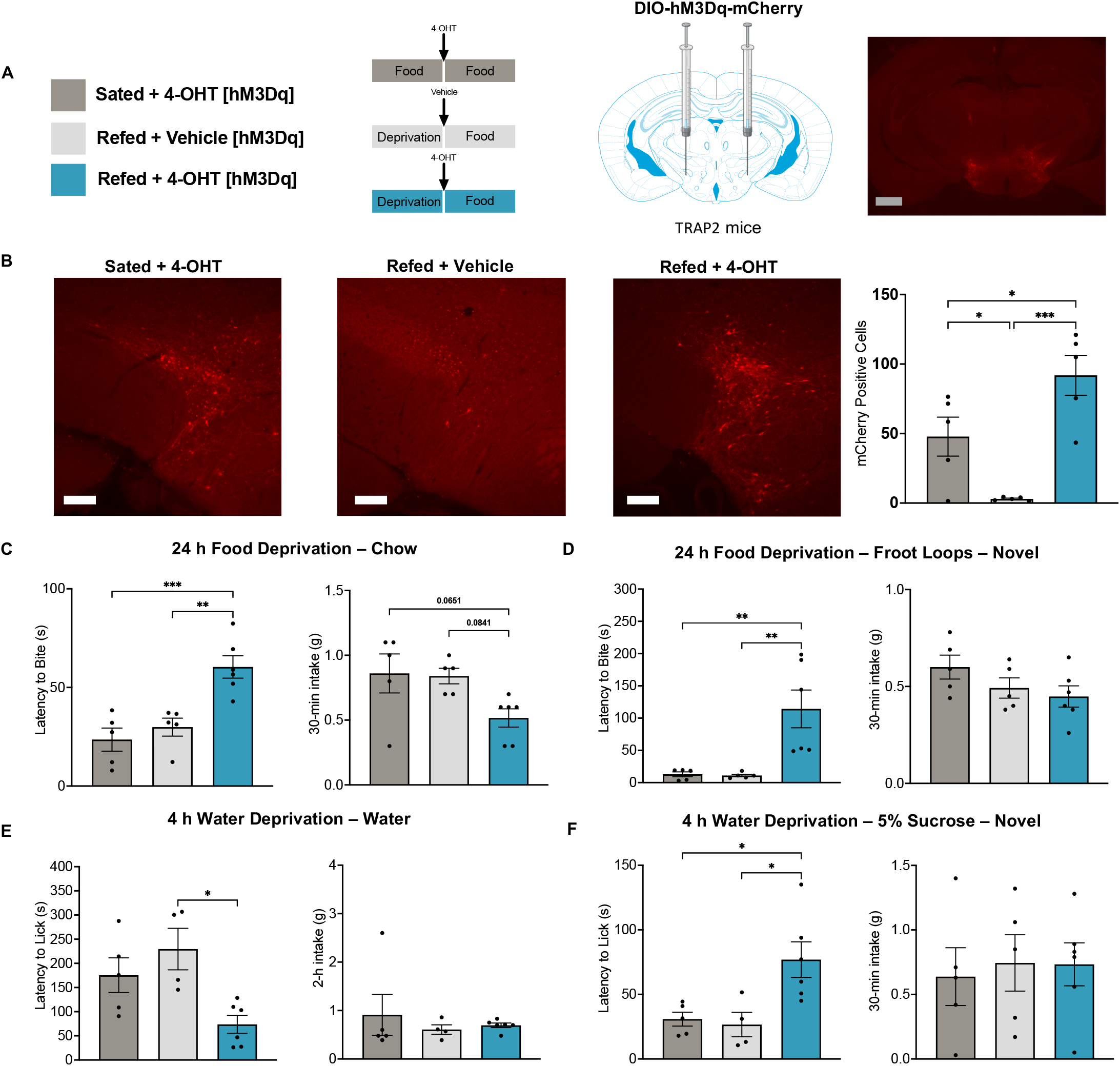
The ensemble of PSTN neurons activated by refeeding delays consumption of familiar and palatable foods. **A**. Experimental design and representative image of mCherry fluorescence showing targeted recombination in the PSTN of a mouse injected with 4-OHT immediately prior to refeeding (scale bars: gray, 500 μm; white, 200 μm). **B**. The number of PSTN mCherry-positive cells illustrates the extent of recombination across experimental conditions. Behavioral testing was performed 30 minutes after injection of the chemogenetic actuator CNO, and 24 hours after food deprivation for chow (**C**) and Froot Loops (**D**), or 4 hours after water deprivation for water (**E**) and sucrose (**F**). In each panel, the latency to initiate feeding/drinking is shown on the left and the amount of food/fluid consumed is shown on the right. Bars represent mean ± s.e.m. and individual values are overlaid. Data were analyzed using one-way ANOVA followed by Tukey’s *posthoc* comparisons when appropriate, *, p<0.05; **, p<0.01; ***, p<0.001.

Mice were again deprived from food for 24 hours and injected with CNO 30 minutes prior regaining access to chow. There was a significant main effect of group on both latency (F_(2,13)_=13.70, p=0.0006) and consumption (F_(2,13)_=4.04, p=0.0433) (**Figure 1C**). Re-activating the refeeding ensemble resulted in significantly longer latencies compared to both sated controls (p=0.0009) and vehicle controls (p=0.0040). Moreover, consumption tended to be reduced compared to sated controls (p=0.0651) and vehicle controls (p=0.0841).

When provided with novel Froot Loops after 24 hours of food deprivation, there was a significant main effect of group on latency (F_(2,13)_=9.77, p=0.0026) but not consumption (F_(2,13)_=1.91, p=0.1871) (**Figure 1D**). Activation of the refeeding ensemble resulted in significantly longer latencies compared to both sated controls (p=0.0066) and vehicle controls (p=0.0058).

We next sought to determine the impact of the PSTN refeeding ensemble on fluid consumption. Mice were deprived of water for 4 hours at the onset of the dark phase and injected with CNO 30 minutes prior to regaining access to water. There was a significant main effect of group on latency (F_(2,12)_=6.55, p=0.0119) but not 2-h consumption (F_(2,12)_=0.36, p=0.7046) (**Figure 1E**). Interestingly, this is the only situation in which activation of a PSTN subpopulation resulted in a significant decrease in latency, here compared to vehicle controls (p=0.0119) and trending for sated controls (p=0.0778). This unique effect may reflect prandial thirst, which is normally triggered during food ingestion to anticipate changes in extracellular fluid osmolality following the absorption of solutes into the bloodstream (4).

When provided with a novel 5% sucrose solution after 4 hours of water deprivation, there was a significant main effect of group on latency (F_(2,12)_=6.82, p=0.0105) but not overall consumption (F_(2,13)_=0.08, p=0.9225) (**Figure 1F**). Activation of the refeeding ensemble resulted in significantly longer latencies compared to sated controls (p=0.0240) and vehicle controls (p=0.0209).

Taken together, these data demonstrate that the PSTN refeeding ensemble is functionally different from the sated, baseline PSTN ensemble and drastically delays feeding initiation in hungry mice but exerts limited control over the amount of food consumed. This conclusion also applies to sucrose ingestion in thirsty mice. Furthermore, the PSTN refeeding ensemble accelerates water drinking. We sought to examine which subpopulation(s) of PSTN neurons might be driving these striking latency effects.

The PSTN contains two non-overlapping populations of neurons expressing either *Tac1* or *Crh*, and both populations respond to refeeding (14). PSTN^*Tac1*^ neurons were previously demonstrated to lower the amount of sucrose consumed by thirsty mice under conditions of novelty (neophobia) or sickness (12). They also contribute to the reduction in meal frequency induced by anorexigenic hormones and reduce food intake upon chemogenetic or optogenetic stimulation (14). In contrast, PSTN^*Crh*^ neurons do not exert such influence (14). These studies, however, did not examine how manipulating the two populations might alter the delay to engage in consummatory behavior. We used Cre-dependent expression of chemogenetic actuators in the PSTN of *Tac1*-Cre and *Crh*-Cre mice to address this question.

### PSTN^*Tac1*^ neurons suppress both the initiation and the execution of consummatory behaviors

Cre-dependent constructs encoding hM4Di, hM3Dq, or mCherry alone were virally transferred into the PSTN of *Tac1*-Cre mice (**Figure 2A**).

**FIGURE 2.**
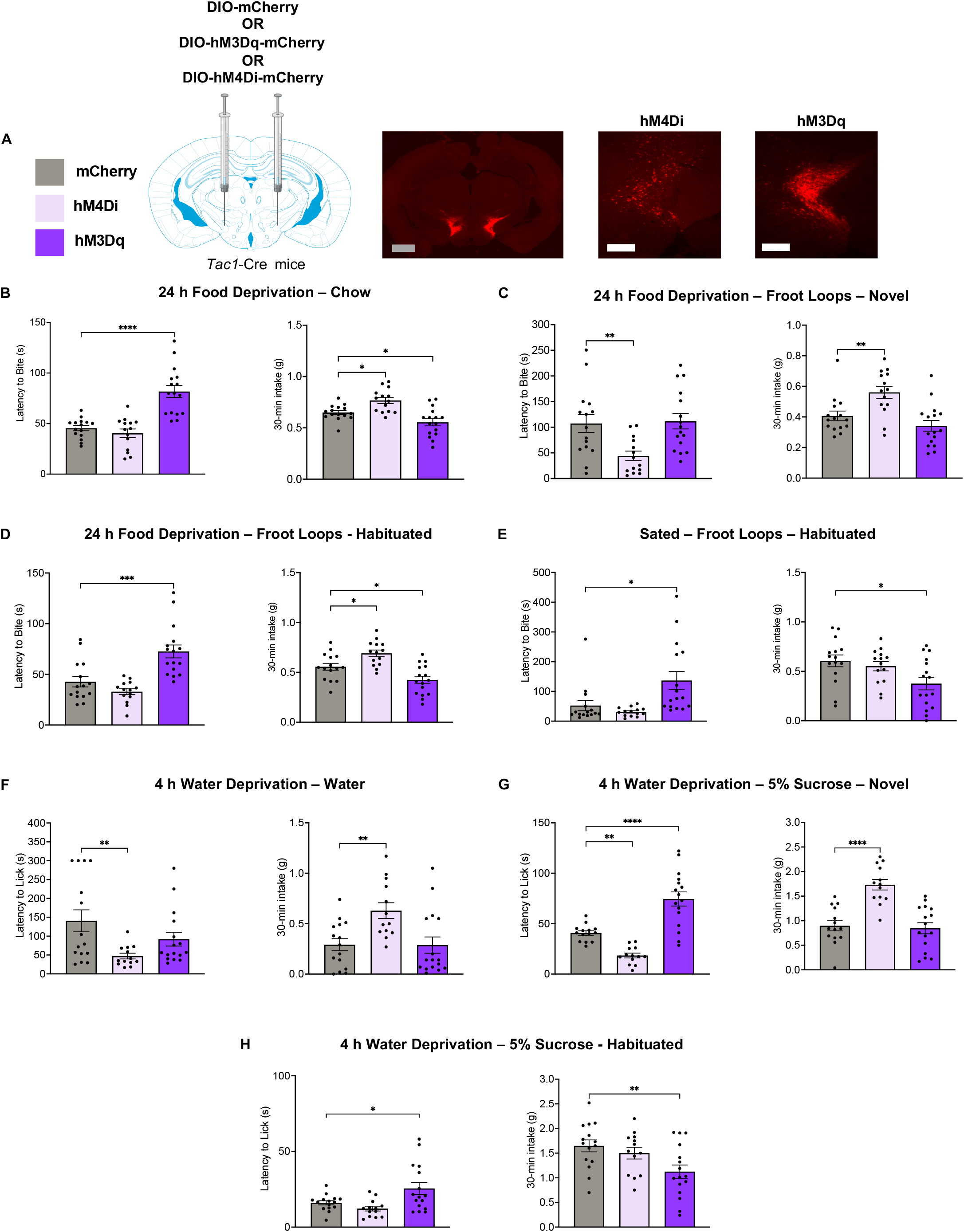
PSTN^*Tac1*^ neurons delay and reduce food and fluid consumption independently of caloric content, metabolic state, and novelty. **A**. Experimental design and representative images of mCherry fluorescence in the PSTN illustrating expression of hM3Dq in PSTN^*Tac1*^ neurons (scale bars: gray, 500 μm; white, 200 μm). Behavioral testing was performed 30 minutes after injection of the chemogenetic actuator CNO, and 24 hours after food deprivation for chow (**B**) and Froot Loops (**C-D**, no food deprivation but same circadian timepoint in **E**), or 4 hours after water deprivation for water (**F**) and sucrose (**G-H**). Testing was performed upon first-time access to Froot Loops and sucrose (**C**, **G**), and again following habituation (**D**, **H**). Froot Loops consumption was also tested after habituation to scheduled access with concurrent *ad libitum* access to chow (**E**). In each panel, the latency to initiate feeding/drinking is shown on the left and the amount of food/fluid consumed is shown on the right. Bars represent mean ± s.e.m. and individual values are overlaid. Data were analyzed using one-way ANOVA followed by Dunnett’s *posthoc* comparisons to mCherry controls, *, p<0.05; **, p<0.01; ***, p<0.001; ****, p<0.0001.

Mice were first assessed for refeeding with familiar chow following 24 hours of food deprivation. There was a significant main effect of vector on both measures (latency: F_2,42_=24.85, p<0.0001; consumption: F_(2,42)_=13.39, p<0.0001) (**Figure 2B**). Activation of PSTN^*Tac1*^ neurons significantly increased the latency to first bite of chow (p<0.0001), while their inhibition had no significant effect (p=0.66). PSTN^*Tac1*^ activation significantly reduced the amount of chow consumed (p=0.0434). In contrast, PSTN^*Tac1*^ inhibition significantly increased chow consumption (p=0.0138). These data indicate that chemogenetic stimulation of PSTN^*Tac1*^ neurons in food-deprived mice is sufficient to delay and reduce chow consumption despite the motivational state produced by hunger. They also show that the endogenous activity of PSTN^*Tac1*^ neurons during refeeding suppresses the consumption of familiar chow, thereby opposing the homeostatic drive to feed.

In a similar experimental design, mice were provided with a novel and presumably palatable high-sugar food, Froot Loops, after 24 hours of food deprivation to assess the sensitivity of PSTN^*Tac1*^ neurons to other dimensions of consumption behaviors. There was a significant main effect of vector on both measures (latency: F_(2,42)_=6.46, p=0.0036; consumption: F_2,42_=9.85, p=0.0003) (**Figure 2C**). Activation did not affect the latency to first bite of Froot Loops (p=0.9670), nor the amount of Froot Loops consumed over 30 min (p=0.3267). In contrast, inhibition resulted in significantly shorter latencies (p=0.0086) and led mice to consume significantly more Froot Loops than controls (p=0.0082). Interestingly, the results here demonstrate that novel palatable foods likely increase the endogenous activity of PSTN^*Tac1*^ neurons to its maximal extent given that no difference was observed in response to chemogenetic activation. This idea is bolstered by the ability of inhibition to reduce latencies to first bite to levels similar to familiar chow and to produce overconsumption compared to controls.

To disentangle the features of Froot Loops being simultaneously a novel and palatable food, mice were given both chow and Froot Loops *ad libitum* for 5 days to dampen novelty. Preference measures for chow versus Froot Loops (no CNO) were collected over a 24-hour period during this habituation period and all mice demonstrated highly significant preference for Froot Loops compared to chow (p<0.0001), thereby confirming the strong palatability of Froot Loops for these mice (**Supplementary Figure S1**). After habituation, mice were again tested with Froot Loops after 24 hours of food deprivation. There was a significant main effect of vector on both measures (latency: F_2,42_=16.18, p<0.0001; consumption: F_2,42_=14.17, p<0.0001) (**Figure 2D**). Activation of PSTN^*Tac1*^ neurons significantly increased latencies (p=0.0003) and reduced intake (p=0.0192) compared to controls. Inhibition did not significantly affect latency (p=0.3238) but led mice to consume significantly more Froot Loops than controls (p=0.0219). Strikingly, manipulations of PSTN^*Tac1*^ neurons while refeeding with habituated Froot Loops yielded effects similar to those observed for familiar chow. In both cases, activation caused a major hesitancy to begin eating and ultimately led to significantly less overall consumption. Inhibition, in contrast, did not increase the speed at which mice took their first bite but did produce overconsumption after 30 minutes. These data also strengthen the idea that PSTN^*Tac1*^ neurons may promote hyponeophagia via increased endogenous activity. Altogether, in hungry mice given access to food, the endogenous activity of PSTN^*Tac1*^ neurons delays feeding initiation selectively under conditions of novelty while it lowers the amount of food consumed regardless of novelty.

In the above-described experiments, food deprivation was used to motivate the mice to readily engage in food consumption. To control for the hunger state as a potential factor underlying our results, consummatory behaviors for habituated Froot Loops were also measured in a sated state, i.e., with *ad libitum* access to chow. Mice were habituated to receiving scheduled access to Froot Loops for 7-8 days to encourage consumption on test day. Upon CNO administration, there was a significant main effect of vector on both latency (F_2,42_=7.16, p=0.0022) and consumption (F_2,42_=4.47, p=0.0174) (**Figure 2E**). Similar to hungry mice given access to familiar food, activation led sated mice to increase their latencies (p=0.0122), while inhibition had no significant effect (p=0.7091). Furthermore, activation significantly decreased consumption (p=0.0128), while no difference was observed in response to inhibition (p=0.7441). These results are compelling given the drastic difference in metabolic state of mice deprived of food for 24 hours versus those in a sated state. These parallel results highlight that a state of hunger is not necessary for PSTN^*Tac1*^ neurons to drive feeding suppression, in line with data published by Kim *et al*. (14).

We next sought to determine the impact of chemogenetic manipulations of PSTN^*Tac1*^ neurons on fluid consumption. For these assays, mice were deprived of water for 4 hours starting at the onset of the dark phase. When access to water was resumed after CNO administration, there was a significant main effect of vector on both measures (latency: F_2,41_=4.65, p=0.0151; consumption: F_2,41_=6.621, p=0.0032) (**Figure 2F**).

Activation did not affect latency to first lick (p=0.1791) nor the amount of water consumed (p=0.9990). Inhibition, however, resulted in significantly shorter latencies (p=0.0078) and significantly higher consumption compared to controls (p=0.0058). These observations reveal that PSTN^*Tac1*^ neurons exert potent inhibitory control over water consumption despite water restriction. Strikingly, the effect of inhibiting PSTN^*Tac1*^ neurons on the latency to rehydrate is comparable to the effect of activating the PSTN refeeding ensemble, which highlights a functional disconnect between these two PSTN subpopulations. This mismatch suggests that only a subset of PSTN^*Tac1*^ neurons may participate in the PSTN refeeding ensemble, in line with the *Fos*/*Tac1* colocalization analysis conducted by Kim *et al*. (14), and that PSTN^*Tac1*^ neurons do not belong to the ensemble driving prandial thirst.

To gain insight into whether PSTN^*Tac1*^ neurons are similarly sensitive to a novel and palatable fluid, mice were provided with a 5% sucrose solution after 4 hours of water deprivation. There was a significant main effect of vector on both measures (latency: F_2,40_=34.58, p<0.0001; consumption: F_2,40_=20.04, p<0.0001) (**Figure 2G**). Activation led to significantly longer latencies when compared to controls, similar to the effect seen with chow and habituated Froot Loops (p<0.0001). In contrast, inhibition resulted in significantly shorter latencies, similar to the effect seen with novel Froot Loops (p=0.0058). After 30 minutes, activation did not alter consumption (p=0.9219), while inhibition led mice to consume significantly more sucrose solution compared to controls (p<0.0001). The results here demonstrate that PSTN^*Tac1*^ neurons similarly modulate the consumption of both palatable solid and liquid foods.

As was done for Froot Loops, mice were habituated to the sucrose solution *ad libitum* for 5 days alongside a separate water bottle to disentangle novelty from the palatable nature of sucrose. Preference measures for water versus 5% sucrose (no CNO) were collected over a 72-hour period and as expected, all mice demonstrated highly significant preference for sucrose compared to water (p<0.0001) (**Supplementary Figure S1**). Upon CNO administration, there was a significant main effect of vector on both measures (latency: F_2,41_=6.36, p=0.0039; consumption: F_2,41_=4.787, p=0.0135) (**Figure 2H**). Parallel to observations with habituated Froot Loops, activation significantly increased latencies (p=0.0292), while no difference was observed with inhibition (p=0.5328). Similarly, activation led mice to consume significantly less habituated sucrose (p=0.0088), while inhibition had no significant effect (p=0.6415). Building on the body of evidence collected in previous appetitive assays, inhibition of PSTN^*Tac1*^ neurons has a stronger overconsumption effect on novel palatable substances than when these items are habituated over time, likely indicating higher endogenous activity of PSTN^*Tac1*^ neurons during hyponeophagia.

### PSTN^*Crh*^ neurons promote the consumption of novel palatable substances

In the PSTN, *Tac1*- and *Crh*-expressing neurons represent non-overlapping populations of the PSTN (14). Accordingly, we tested whether PSTN^*Crh*^ neurons might exert a differential influence on consummatory behaviors compared to the PSTN^*Tac1*^ population. A Cre-dependent construct encoding the excitatory designer receptor hM3Dq was virally transferred into the PSTN of *Crh*-Cre mice (**Figure 3A**).

**FIGURE 3.**
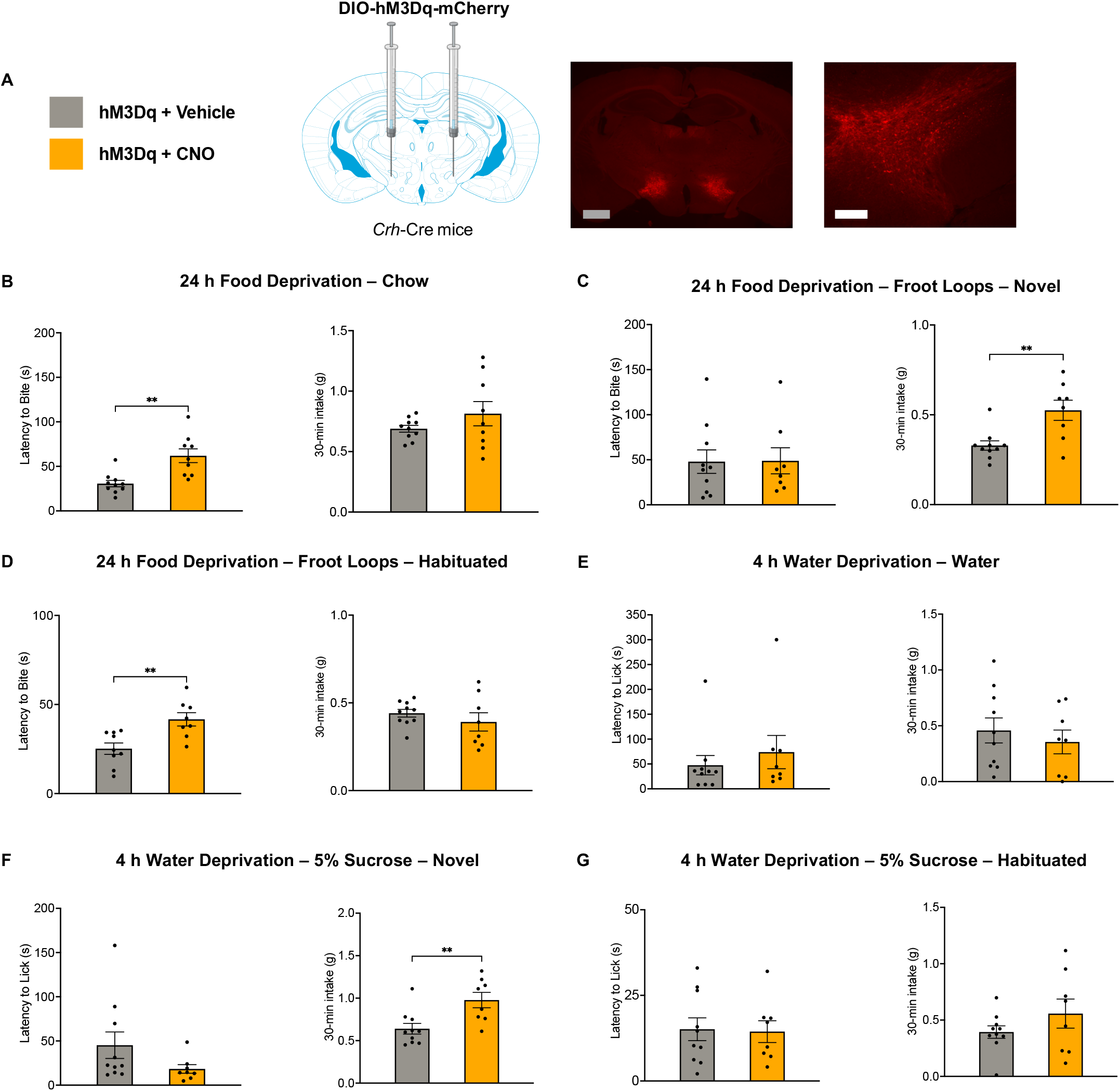
PSTN^*Crh*^ neurons delay refeeding of familiar foods and increase consumption of novel palatable substances. **A**. Experimental design and representative images of mCherry fluorescence illustrating expression of hM3Dq in PSTN^*Crh*^ neurons (scale bars: gray, 500 μm; white, 200 μm). Behavioral testing was performed 30 minutes after injection of the chemogenetic actuator CNO or vehicle, and 24 hours after food deprivation for chow (**B**) and Froot Loops (**C-D**), or 4 hours after water deprivation for water (**E**) and sucrose (**F-G**). Testing was performed upon first-time access to Froot Loops and sucrose (**C**, **F**), and again following habituation (**D**, **G**). In each panel, the latency to initiate feeding/drinking is shown on the left and the amount of food/fluid consumed is shown on the right. Bars represent mean ± s.e.m. and individual values are overlaid. Data were analyzed using unpaired t tests, *, p<0.05; **, p<0.01.

Mice were deprived from food for 24 hours and injected with CNO or vehicle 30 minutes prior to the beginning of experimental sessions. Mice were first assessed for refeeding with familiar chow (**Figure 3B**). Activation of PSTN^*Crh*^ neurons by CNO resulted in significantly longer latencies compared to vehicle (t_18_=3.15, p=0.0055). The amount of chow consumed in 30 minutes, however, was indistinguishable between treatments (t_18_=0.12, p=0.9101). The results demonstrate that PSTN^*Crh*^ neurons can drive hesitancy to first bite of chow, but do not influence overall consumption.

Mice were provided with novel Froot Loops after 24 hours of food deprivation to assess the influence of PSTN^*Crh*^ neurons on other dimensions of appetitive behaviors. Activation of PSTN^*Crh*^ neurons did not affect the latency to first bite (t_16_=0.05, p=0.9627) but significantly increased Froot Loops consumption (t_16_=3.39, p=0.0037) (**Figure 3C**). These results demonstrate that activation PSTN^*Crh*^ neurons leads to overconsumption of a novel, palatable food, i.e., similar to the effect of inhibiting PSTN^*Tac1*^ neurons. Altogether, these observations suggest that the two populations oppose each other’s function in situations of hyponeophagia.

To disentangle the novelty of Froot Loops from their palatability, mice were given both chow and Froot Loops *ad libitum* for 5 days. After habituation, mice were again tested for Froot Loops after 24 hours of food deprivation. As was observed with chow, activation resulted in significantly longer latencies (t_15_=3.37, p=0.0042), with no significant effect on consumption (t_16_=0.95, p=0.3574) (**Figure 3D**).

We next sought to determine the impact of chemogenetic excitation of PSTN^*Crh*^ neurons on liquid consumption after 4 hours of water deprivation. When access to water was resumed after CNO or vehicle administration, there was no significant effect of treatment on latency (t_16_=0.72, p=0.4850) or consumption (t_16_=0.65, p=0.5231) (**Figure 3E**), indicating that *Crh* neurons do not participate in the PSTN circuit triggering prandial thirst.

When provided with a novel, palatable 5% sucrose solution, there was no significant effect of treatment on latency (t_16_=1.54, p=0.1432) but CNO increased sucrose consumption compared to vehicle (t_16_=3.15, p=0.0062) (**Figure 3F**). These results are congruent with the first-time Froot Loops assay, such that excitation of PSTN^*Crh*^ neurons significantly increase the consumption of novel, palatable solid and liquid substances without affecting the motivation to begin consuming.

Mice were habituated to the sucrose solution *ad libitum* for 5 days to disentangle novelty from the palatable nature of sucrose. There was no significant main effect of treatment on latency (t_16_=0.15, p=0.8853) or consumption (t_16_=1.26, p=0.2276) (**Figure 3G**). Altogether, activation of PSTN^*Crh*^ neurons did not affect the consumption of familiar chow, habituated Froot Loops, or habituated sucrose and selectively increased first-time consumption of Froot Loops and sucrose. These results underscore the sufficiency of PSTN^*Crh*^ neurons to promote the consumption of novel palatable substances without altering the latency to initiate their ingestion, a pattern that does not support the participation of *Crh* neurons in the PSTN refeeding ensemble.

### PSTN neurons projecting to the CeA suppress feeding initiation

A dual-vector, pathway-specific chemogenetic approach was employed to determine the contribution of PSTN projection neurons. Previous neuroanatomic tracing demonstrated prominent projections from the PSTN to the CeA and to the BNST (7, 14, 15), two structures that are involved in feeding regulation (31–33). While PSTN^*Tac1*^ neurons send inputs to both the CeA and the BNST, food consumption is reduced by photoactivation of CeA-projecting, but not BNST-projecting, PSTN^*Tac1*^ neurons, highlighting the differential role of these two pathways (14). We therefore sought to assess the influence of CeA- and BNST-projecting PSTN neurons in the control of feeding initiation. In C57BL/6J mice, a retrograde Cre-expressing virus was injected into the CeA or the BNST permitting the Cre-dependent expression of hM3Dq and KORD viral constructs co-injected into the PSTN to excite or inhibit those projections neurons upon administration of CNO or SalB, respectively (**Figure 4A**).

**FIGURE 4.**
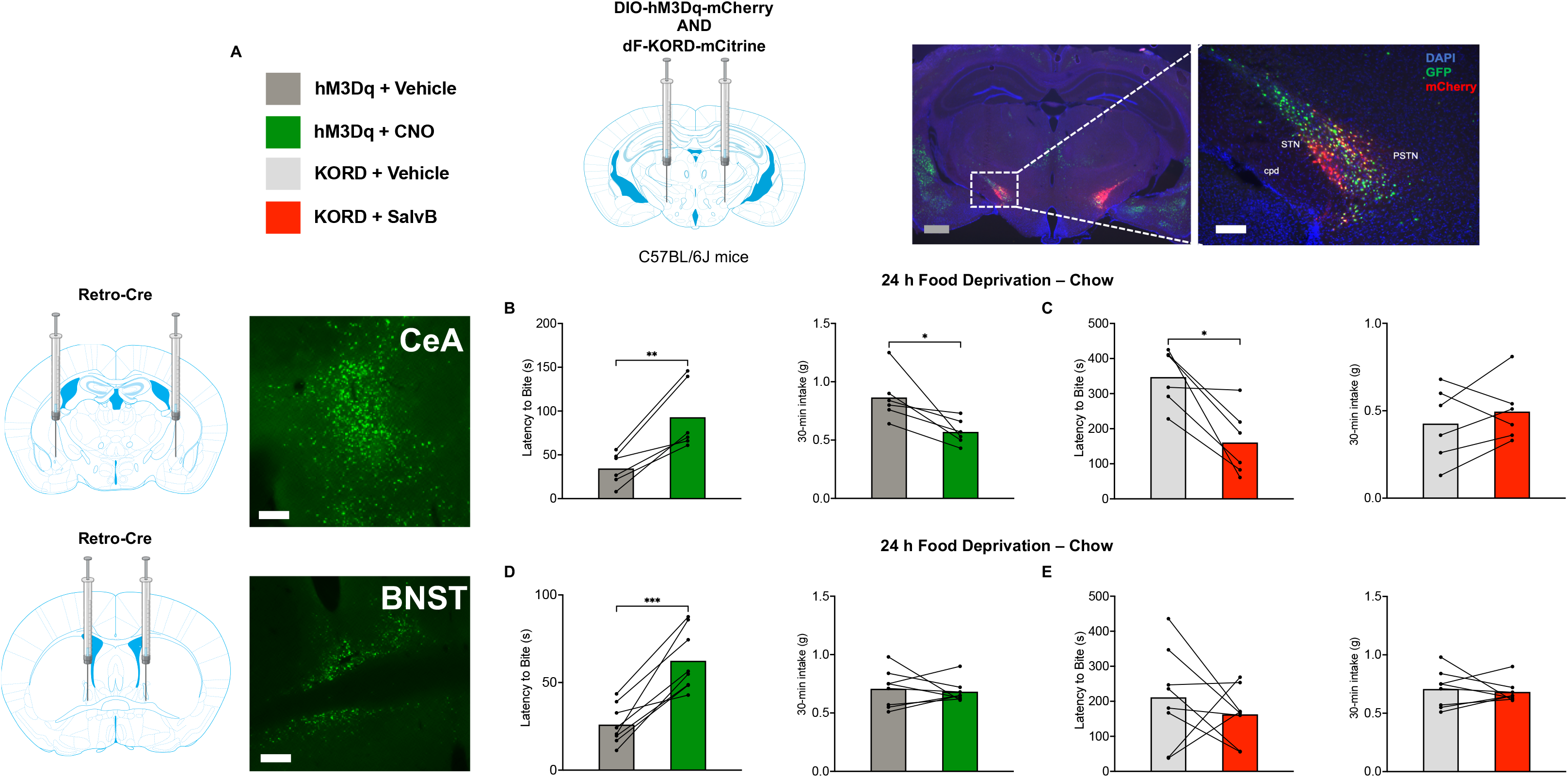
CeA-but not BNST-projecting PSTN neurons mediate refeeding delay. **A**. Experimental design and representative images of GFP fluorescence illustrating Cre expression and mCherry fluorescence illustrating hM3Dq expression in the PSTN (scale bars: gray, 500 μm; white, 200 μm). Cre was expressed from a retrograde vector injected either in the CeA (**B-C**) or in the BNST (**D-E**). Behavioral testing was performed 30 minutes after injection of the chemogenetic actuators CNO (**B**, **D**) or SalB (**C**, **E**), or their respective vehicles, 24 hours after food deprivation. Chow was presented either in the home cage (**B**, **D**) or in a novel arena (**C**, **E**). In each panel, the latency to initiate feeding/drinking is shown on the left and the amount of food/fluid consumed is shown on the right. Bars represent mean ± s.e.m. and individual values are overlaid. Data were analyzed using paired t tests, *, p<0.05; **, p<0.01; ***, p<0.001.

Mice were deprived from food for 24 hours and injected with chemogenetic ligands (CNO, SalB, or their respective vehicles) 30 minutes prior to refeeding with familiar chow. SalB testing was conducted in an open arena to increase feeding latency and facilitate the detection of downward shifts. PSTN^CeA^ activation significantly increased latency (t_5_=4.92, p=0.0044) and reduced consumption (t_5_=3.09, p=0.0272) (**Figure 4B**). Conversely, PSTN^CeA^ inhibition significantly shortened latency (t_5_=3.97, p=0.0106) but did not affect consumption (t_5_=0.90, p=0.4118) (**Figure 4C**). PSTN^BNST^ activation also increased latency (t_7_=6.57, p=0.0003) but did not alter consumption (t_7_=0.40, p=0.6994) (**Figure 4D**). PSTN^BNST^ inhibition had no significant effect on either latency (t_7_=0.82, p=0.4419) or consumption (t_7_=0.40, p=0.6994) (**Figure 4E**). Altogether, these data point to a prominent role of the endogenous activity of PSTN^CeA^ neurons in delaying food consumption initiation in hungry mice.

### CeA-projecting PSTN^*Tac1*^ neurons delay refeeding

Given that both PSTN^*Tac1*^ and PSTN^*Crh*^ neurons can delay feeding initiation in food-deprived mice given access to chow (**Figures 2B** and **3B**) and that they both project to the CeA (12, 14), either cell type may contribute to the influence of the PSTN^CeA^ projection on refeeding latency. To explore this possibility, we used a dual-vector, intersectional strategy to drive the expression of hM3Dq in PSTN^*Tac1*^ and PSTN^*Crh*^ projecting to the CeA. This selective targeting was driven by the expression of Cre in *Tac1*-Cre or *Crh*-Cre mice respectively, the injection of a retrograde Flp-encoding vector in the CeA, and the injection of an INTRSECT (INTronic Recombinase Sites Enabling Combinatorial Targeting (27)) vector encoding hM3Dq-mCherry upon Cre AND Flp recombination (**Figure 5A**) in the PSTN.

**FIGURE 5.**
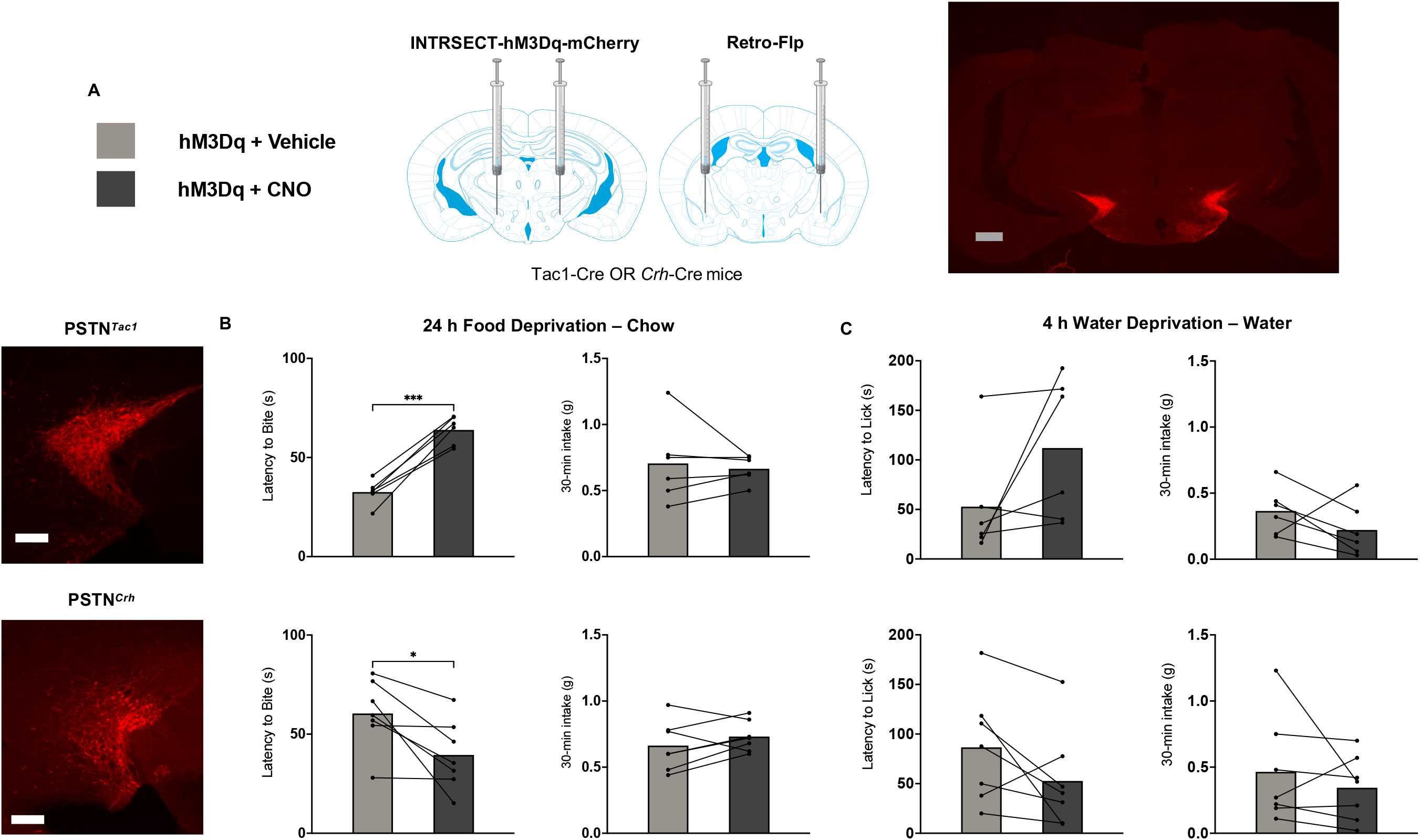
CeA-projecting PSTN^*Tac1*^ neurons delay refeeding while CeA-projecting PSTN^*Crh*^ neurons hasten refeeding. **A**. Experimental design and representative images of mCherry fluorescence illustrating hM3Dq expression in CeA-projecting PSTN^*Tac1*^ and PSTN^*Crh*^ neurons (scale bars: gray, 500 μm; white, 200 μm). Behavioral testing was performed 30 minutes after injection of the CNO or vehicle, and 24 hours after food deprivation (**B**) or 4 hours after water deprivation (**C**). In each panel, the latency to initiate feeding/drinking is shown on the left and the amount of food/fluid consumed is shown on the right. Bars represent mean ± s.e.m. and individual values are overlaid. Data were analyzed using paired t tests, *, p<0.05; ***, p<0.001.

Following 24-hour food deprivation, activating CeA-projecting PSTN^*Tac1*^ significantly increased the latency to initiate familiar chow consumption (t_5_=9.32, p=0.0002) but did not affect the amount consumed (t_5_=0.434, p=0.6822) (**Figure 5B**). These results further underscore the importance of this molecularly defined projection given that activation of PSTN^*Tac1*^ neurons (**Figure 2B**) and CeA-projecting neurons (**Figure 4B**), in isolation or combined, significantly increases latency to first bite of familiar chow. However, total consumption of chow was not affected by the activation of CeA-projecting PSTN^*Tac1*^ neurons, in contrast to the significant decreases observed when either PSTN^*Tac1*^ neurons (**Figure 2B**) or CeA-projecting neurons (**Figure 4B**) were activated independently. Most relevant to our goal of characterizing the neuronal subpopulations activated by food access in hungry mice, this pattern is identical to the effect of re-activating the PSTN refeeding ensemble (**Figure 1C**).

We then determined the effect of CeA-projecting PSTN^*Tac1*^ activation on water drinking. When access to water was resumed after CNO or vehicle administration, there was no significant effect of treatment on latency (t_5_=1.82, p=0.1270) or consumption of water (t_5_=1.32, p=0.2431) (**Figure 5C**), indicating that these neurons do not contribute to the PSTN subpopulation triggering prandial thirst.

Activating CeA-projecting PSTN^*Crh*^ after 24-hour food deprivation significantly reduced the latency to initiate familiar chow consumption (t_6_=3.07, p=0.0218) without affecting the amount consumed (t_6_=1.308, p=0.2388) (**Figure 5D**). CeA-projecting PSTN^*Crh*^ thereby stands out as the only population in our study sufficient to hasten refeeding, which underscores not only an opposing role of *Tac1* and *Crh* subsets within the PSTN^CeA^ projection, but also the unique influence of CeA-projecting *Crh* cells among all PSTN^*Crh*^ neurons (**Figure 3B**).

Activating CeA-projecting PSTN^*Crh*^ neurons after 4 hours of water deprivation had no significant effect on the latency to initiate drinking (t_6_=1.94, p=0.1011) or the amount of water consumed (t_6_=0.914, p=0.3959) (**Figure 5E**), similar to the results observed with general PSTN^*Crh*^ activation (**Figure 3E**).

## Discussion

The results of this study demonstrate that the PSTN exerts a profound effect on the motivation to initiate feeding. The PSTN is known to become active in response to diverse anorexigenic signals including binge-like refeeding following food deprivation, sickness, nutrient-deficient diets, novelty, and homeostatic hormonal signaling (8–14). Our study highlights that the functional significance of this activation is to suppress the initiation of food consumption, an influence that extends to palatable solids and calorie-containing liquids, with minimal impact on the amount of food consumed (**Figure 1**). We further find that these refeeding-sensitive cells are phenotypically similar to PSTN^*Tac1*^ neurons (**Figure 2**) but distinct from PSTN^*Crh*^ neurons (**Figure 3**). Furthermore, we show that PSTN^CeA^ neurons activated by food access in a hungry state play a prominent role in suppressing feeding initiation (**Figure 4**). Finally, specifically activating CeA-projecting PSTN^*Tac1*^ neurons was sufficient to delay refeeding, while activating CeA-projecting PSTN^*Crh*^ neurons hastened refeeding, highlighting the functional heterogeneity of PSTN subpopulations within a given anatomical pathway (**Figure 5**). Altogether, our data suggests that CeA-projecting PSTN^*Tac1*^ neurons represent a major component of the PSTN ensemble that drives hesitancy in hungry animals given access to food. By enhancing our understanding of the brain mechanisms controlling feeding onset, independently of consumption *per se*, this finding may have therapeutic relevance for the treatment of eating disorders. Activating CeA-projecting PSTN^*Tac1*^ neurons could help alleviate the craving component of bulimia nervosa and binge-eating disorders, while inhibiting this circuit may loosen pathological self-control in restrictive anorexia nervosa – and neither of these manipulations would be expected to negatively affect homeostatic food consumption. Future studies will aim to elucidate the molecular signaling events underlying the suppression of feeding initiation by CeA-projecting PSTN^*Tac1*^ neurons to afford pharmacological access to this circuit.

While PSTN neurons activated upon refeeding of hungry animals with familiar chow had a minimal impact on the amount of food consumed, our results from chemogenetic inhibition show that the endogenous activity of PSTN^*Tac1*^ neurons exerts a strong influence on consumption when mice are given access to a novel palatable food or fluid, consistent with the effect reported by Barbier *et al*. in mice given access to sucrose for the first time (12). We found that this influence is reduced by habituation, which is consistent with reduced c-Fos induction in the PSTN of rats habituated to sucrose consumption (12), and further blunted by removing the state of metabolic need triggered by food deprivation. As expected, reducing the endogenous tone of PSTN^*Tac1*^ activity by using a familiar food or fluid was necessary for chemogenetic stimulation to significantly reduce consumption, in line with the effect of optogenetic stimulation of PSTN^*Tac1*^ neurons in *ad libitum* fed mice (14). Interestingly, the endogenous activity of PSTN^*Tac1*^ neurons also suppressed water drinking, indicating that the influence of this subpopulation on consummatory behavior is independent of caloric content. This observation contrasts starkly with the influence of the PSTN refeeding ensemble on water intake (discussed in more detail below), indicating that these two PSTN subpopulations only have a partial overlap.

Another important outcome of our study was the opposing influence of PSTN^*Tac1*^ vs. PSTN^*Crh*^ neurons on palatable food consumption, which was suppressed by the former but increased by the latter. While previous studies had already identified an important role of PSTN^*Tac1*^ in feeding suppression (12, 14), less is known regarding the physiological role of PSTN^*Crh*^ neurons (7). PSTN^*Crh*^ neurons are activated by refeeding (14) and *Crh* expression in the PSTN is upregulated by an anorexia-inducing valine-deficient diet (9), but they do not influence food consumption (14 and results reported here). Aside from consummatory behaviors, PSTN^*Crh*^ neurons have been shown to mediate defensive responses to acute predator threats and to regulate REM sleep (34).

Our observation that their activation promotes the consumption of sweet food and liquids suggests an important role in the control of hedonic feeding. Future studies will be needed to determine if this influence may extend to the consumption of other rewarding substances (e.g., addictive drugs) or experiences (e.g., social interaction).

Our results also uncover an unexpected implication of the PSTN in water drinking initiation. Re-activating the PSTN refeeding ensemble accelerated the onset of water consumption, raising the possibility that the PSTN may be connected to circuits controlling prandial thirst, such as subfornical organ *Nos1* neurons or posterior pituitary-projecting, vasopressin-secreting neurons (4, 35-37). While additional experiments will be needed to probe the anatomical and functional validity of this putative connection, our findings suggest that it does not involve PSTN *Tac1* nor *Crh* neurons. Interestingly, the shortening of drinking latency driven by the PSTN refeeding ensemble was selective for water, as an opposite effect was observed when mice were given access to a sucrose solution. Since latency modulation occurs before the mice get to sample the fluid’s taste, the differential effects of the PSTN refeeding ensemble on water vs. sucrose drinking initiation probably involve olfactory cues that enable the mice to predict the composition of the available solution (38).

In conclusion, our study identifies a novel circuit that suppresses feeding initiation despite hunger, which emphasizes the relevance of the PSTN as a key brain site in the control of appetite and food rejection (18) and has important implications for our understanding of the neural mechanisms that may be disrupted in eating disorders.

## Acknowledgments

We wish to thank Maya Mehta and Kera Norris for their assistance with histological analysis.

## Competing interests

The authors have no conflicts of interest to disclose.

## CRediT authorship contribution statement

**Jeffery L Dunning**: Conceptualization, Investigation, Methodology, Formal analysis, Writing – original draft. **Catherine Lopez**: Investigation. **Colton Krull**: Investigation. **Max Kreifeldt**: Investigation. **Maggie Angelo**: Investigation. **Charu Ramakrishnan**: Methodology, Resources, Validation. **Karl Deisseroth**: Methodology, Resources, Validation, Supervision. **Candice Contet**: Conceptualization, Funding acquisition, Project administration, Supervision, Writing – review & editing.

## Funding

This work was supported by National Institutes of Health research grants AA026685, AA027636, AA006420, and AA027372, as well as training grant AA007456. These funding sources were not involved in study design, data collection, analysis, or interpretation, nor decision to publish.

---

**SUPPLEMENTAL FIGURE 1.**
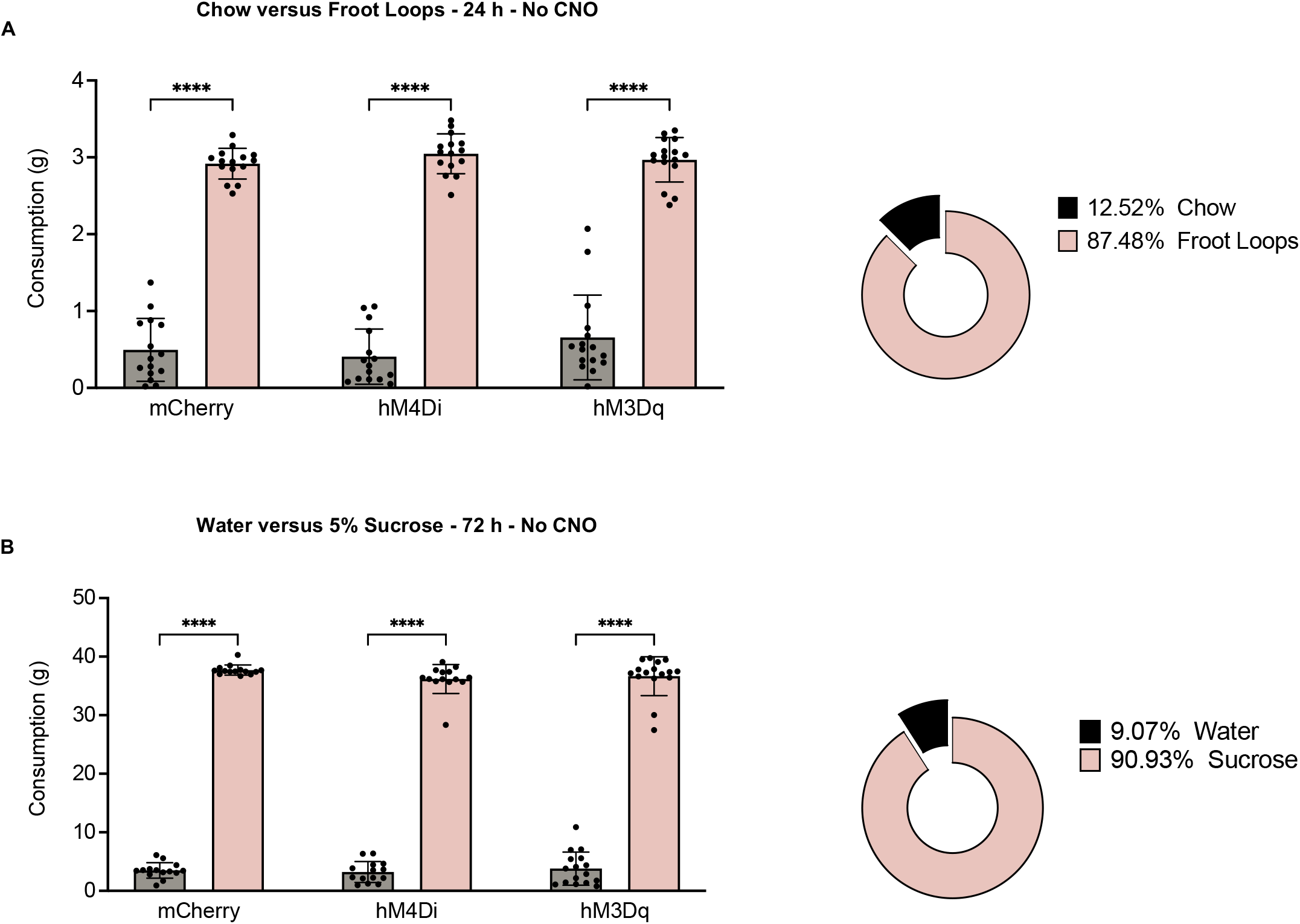
Mice demonstrate robust preference for Froot Loops and sucrose. To interpret experimental data obtained upon chemogenetic manipulation, all mice were given the choice to consume (**A**) regular chow and Froot Loops during a 24-hour period, or (**B**) water and a 5% sucrose solution during a 72-hour period. No CNO was administered. Consumption is measured in grams on the left and average percent of total consumption is illustrated on the right. Data were analyzed using two-way ANOVA followed by Tukey’s *posthoc* comparisons, ****, p<0.0001.

## Notes

### Competing Interest Statement

The authors have declared no competing interest.

